# The COP9 Signalosome Suppresses Cardiomyocyte Necroptosis

**DOI:** 10.1101/2019.12.19.883322

**Authors:** Peng Xiao, Changhua Wang, Megan T. Lewno, Penglong Wu, Jie Li, Huabo Su, Jack O. Sternburg, Jinbao Liu, Xuejun Wang

**Author notes:** These authors contributed equally (Peng Xiao, Changhua Wang). Address correspondence to: Dr. Xuejun Wang, Division of Basic Biomedical Sciences, Sanford School of Medicine of the University of South Dakota, 414 East Clark Street, Vermillion, SD 57069, USA.

## Abstract

**Background:** Loss of cardiomyocyte (CMs) due to apoptosis and regulated necrosis contributes to heart failure. However, the molecular mechanisms governing regulated CM necrosis remain obscure. The COP9 signalosome (CSN) formed by 8 unique protein subunits (COPS1 through COPS8) functions to deneddylate Cullin-RING ligases (CRLs), thereby regulating the functioning of the CRLs. Mice with CM-restricted knockout of *Cops8* (Cops8-cko) die prematurely, following reduced myocardial performance of autophagy and the ubiquitin-proteasome system (UPS) as well as massive CM necrosis. This study was aimed to determine the nature and underlying mechanisms of the CM necrosis in Cops8-cko mice.

**Methods:** We examined myocardial expression and activities of key proteins that reflect the status of the RIPK1-RIPK3 pathway, redox, and caspase 8 in Cops8-cko mice. Moreover, we used in vivo CM uptake of Evan’s blue dye (EBD) as an indicator of necrosis and performed Kaplan-Meier survival analyses to test whether treatment with a RIPK1 kinase inhibitor (necrostatin-1) or an antioxidant (N-acetyl-L-cysteine), global knockout of the *RIPK3* or the *Ppif* gene, CM-restricted knockout of the *Nrf2* gene, or cardiac *HMOX1* overexpression could rescue the Cops8-cko phenotype.

**Results:** Compared with littermate control mice, myocardial protein levels of RIPK1, RIPK3, MLKL, the RIPK1-bound RIPK3, protein carbonyls, full-length caspase 8, Nrf2, Ser40-phosphorylated Nrf2 and BCL2, as well as histochemical staining of superoxide anions were significantly increased but the cleaved caspase 8 and the overall caspase 8 activity were markedly decreased in Cops8-cko mice, indicating that the RIPK1-RIPK3 and the Nrf2 pathways are activated and caspase 8 activation is suppressed by Cops8-cko. Continuous necrostatin-1 infusion initiated at 2 weeks of age nearly completely blocked CM necrosis at 3 weeks and markedly delayed premature death of Cops8-cko mice. *RIPK3* haploinsufficiency or cardiac-specific *Nrf2* heterozygous knockout discernably attenuated CM necrosis and/or delayed mouse premature death; conversely, *Ppif* knockout, N-acetyl-L-cysteine treatment, and cardiac overexpression of HMOX1 exacerbated CM necrosis and mouse premature death.

**Conclusions:** Cardiac Cops8/CSN malfunction causes RIPK1-RIPK3 mediated CM necroptosis in mice; sustained Nrf2 activation and reductive stress pivot cardiomyocytes to necroptosis when autophagy and the UPS are impaired; and the CSN plays an indispensable role in suppressing CM necroptosis.

## Introduction

The COP9 signalosome (CSN) is a highly conserved protein complex formed by 8 unique protein subunits (COPS1 through COPS8). The known biochemical activity of the CSN is to serve as the deneddylase to remove NEDD8 from a neddylated cullin in the Cullin-RING ligase complexes (CRLs) via a process known as deneddylation.^1^ The catalytic center of the CSN is harbored in COPS5 but COPS5 exerts proper deneddylating activity only when it is incorporated into the CSN holocomplex formed by all 8 subunits;^2^ hence, loss of any of the COPS subunits will impair Cullin deneddylation. Cullin functions as a scaffold in CRLs which are the largest family of ubiquitin E3s and, by estimate, responsible for the ubiquitin-dependent degradation of approximately 20% of cellular proteins.^3^ It has been suggested that CRLs play an important role in the degradation of misfolded proteins in the heart.^4^ The Skp1-Cul1-F-box (SCF) E3s are the prototype of CRLs and classified as the CRL1 class. There are at least 7 other classes of CRLs.^5^ Cullin neddylation and deneddylation regulate the cyclic assembly and disassembly of CRLs, which is essential for remodeling CRLs to meet timely the need to ubiquitinate specific substrate proteins within the cell.^6^ Thus the CSN by virtue of Cullin deneddylation plays an indispensable role in regulating the ubiquitination of a significant proportion of cellular proteins. We have previously reported that cardiomyocyte (CM)-restricted knockout (KO) of the *Cops8* gene (Cops8^CKO^) in mice initiated at the perinatal period leads to massive CM necrosis, dilated cardiomyopathy, and mouse premature death, which is preceded by perturbation of not only the ubiquitin-proteasome system (UPS) but also the autophagic-lysosomal pathway (ALP).^7, 8^ Similar findings were also observed in mice with adult-onset Cops8^CKO^.^9^ The present study was performed to investigate why Cops8 deficiency in CMs causes necrosis.

Morphologically, cell death can be generally classified into necrosis (AKA, lytic cell death) and apoptosis (AKA, non-lytic cell death).^10^ Necrosis is featured by the loss of cell membrane integrity, which allows free entry of extracellular fluid into the cell. This process leads to cell swelling, rupturing, and subsequent releasing of cellular contents into the extracellular space; hence, necrosis will inevitably trigger inflammation. Conversely, apoptosis is a well-known and well-characterized form of programmed or regulated cell death that requires caspase activation via either the mitochondrial or the extrinsic pathway. When a cell undergoes apoptosis in a tissue, the cell keeps its membrane sealed well and, even at the late stage, the apoptotic cell breaks into smaller pieces known as apoptotic bodies, each of which is capsuled by membrane. Hence, apoptosis generally does not trigger inflammation and is a much cleaner form of cell death than necrosis.^11^ Recent advances in cell death research have further unveiled that a significant portion of necrosis can also be regulated cell death, known as regulated necrosis, of which death receptor-triggered necrosis is known as necroptosis.^11^ Originally identified in caspase 8 deficient or inhibited cells, the induction of necroptosis by TNFα is now known to require the formation of necrosomes consisting of receptor interacting protein kinase 1 (RIPK1), RIPK3, and a pseudo-kinase termed mixed lineage kinase-like protein (MLKL). In the canonical pathway by which the activation of TNFα receptor 1 (TNFR1) induces necroptosis, the kinase activities of both RIPK1 and RIPK3 are required to phosphorylate MLKL. Phosphorylated MLKL forms amyloid-like oligomers, which will then translocate and incorporate into the plasma membrane; ultimately, producing pores on the membrane which will lead to the cell swelling and plasma membrane rupture.^11^ Ubiquitination plays an essential role in the regulation of both the kinase activity of RIPK1 and the activation of caspase 8. For example, in TNFR1 signaling, both K63-linked and methionine 1 linear ubiquitination of RIPK1 are required for the incorporation of RIPK1 into the complex 1 and thereby promote NFκB activation and cell survival,^12, 13^ whereas K48-linked polyubiquitination of RIPK1 mediates its proteasomal degradation.^14, 15^ Cullin3 (Cul3)-based polyubiquitination of caspase 8 drives full activation and processing of caspase 8, which leads to activation of the extrinsic apoptotic pathway.^16^ However, it remains unclear how the malfunction of the CSN, a major regulator of CRLs, impacts these cell death pathways although ablation of various *Cops* genes and the chemical inhibition of the CSN are known to induce cell death.^7–9, 17, 18^

Loss of the cardiomyocyte (CM) as a result of apoptosis and/or various forms of regulated necrosis contributes to heart failure,^11, 19^ a leading cause of disability and death in humans. Findings from analyzing biochemical markers of necroptosis in the myocardium of humans with end-stage heart failure resulting from myocardial infarction (MI) or dilated cardiomyopathy indicate an involvement of necroptosis in the development of heart failure.^20^ A genetic variant in the *RIPK3* promoter region associated with increased *RIPK3* transcription may contribute to the poor prognosis of heart failure patients.^21^ Animal experiments demonstrated an important role for necroptosis in post-MI remodeling,^22^ myocardial ischemia/reperfusion (I/R) injury, cardiotoxicity of doxorubicin treatment,^23, 24^ and paraquat-induced cardiac contractile dysfunction.^25^ Mechanistically, one elegant study has shown that cardiac necroptosis induced by I/R injury or doxorubicin treatment requires RIPK3 but not RIPK1 and MLKL; the upregulated RIPK3 phosphorylates and activates the calcium/calmodulin-dependent protein kinase II (CaMKII) and thereby opens mitochondrial permeability transition pore (MPT) to induce CM necroptosis.^23^ However, more recent evidence suggests that the RIPK3-MLKL axis may still be important for myocardial necroptosis during I/R injury.^24^ Myocardial I/R was shown to induce myocardial dysregulation of both strands (5p and 3p) of miR-223 in mice and this dysregulation induces cardiac necroptosis during I/R by acting on TNFR1 and other points upstream of RIPK3.^26^ Consistent with the crucial role of transforming growth factor beta-activated kinase 1 (TAK1) and TNFR-associated protein 2 (TRAF2) in TNFR1-triggered survival signaling, CM-restricted ablation of the gene encoding TAK1 or TRAF2 in mice causes CM apoptosis and necroptosis and thereby increases the propensity for heart failure.^27, 28^ Taken together, these studies strongly support the proposition that CM necroptosis plays an important role in the development of heart failure from common etiologies such as ischemic heart disease, dilated cardiomyopathy, and perhaps hypertensive heart disease. Therefore, a better understanding of the molecular mechanisms governing CM necroptosis may provide new therapeutic strategies to prevent or more effectively treat heart failure.

The present study determined the nature and underlying mechanisms of the CM necrosis observed in Cops8^CKO^ mice. It revealed that CM necrosis induced by Cops8 deficiency or CSN impairment was associated with increased interaction of RIPK1 with RIPK3, decreases in caspase 8 activation, and sustained activation of the Nrf2-BCL2 pathway. Moreover, inhibition of RIPK1 kinase activity and the haploinsufficiency of either RIPK3 or Nrf2, but not ablation of the gene encoding Cyclophilin D or augmentation of the antioxidant capacity, were able to significantly attenuate Cops8^CKO^-induced CM necrosis and delay mouse premature death. Hence, this study demonstrates that COPS8 deficiency or CSN impairment causes CM necroptosis in mice through activating the RIPK1-RIPK3 pathway, sustaining Nrf2 activation and impairing caspase 8 activation, which establishes Cops8/the CSN as a crucial suppressor of CM necroptosis and unravels novel mechanisms for cardiac UPS and ALP malfunction in injuring the heart. To our knowledge, this study also provides the first demonstration that sustained Nrf2 activation and reductive stress can steer cardiomyocytes to necroptosis when autophagy and the UPS are malfunctioned, a combination that is frequently implicated in human heart disease.

## Materials and Methods

### Animal models

Perinatal cardiomyocyte-restricted ablation of the *Cops8* gene (Cops8^CKO^) was achieved in C57BL/6J inbred mice as we previously reported.^7^ The creation of RIPK3 null mice was previously described.^29^ Mice with germline knockout of the *Ppif* gene (encoding Cyclophilin D) were provided by Dr. Jeffrey Molkentin of University of Cincinnati.^30^ The floxed mutant mice harboring *loxP* sites flanking exon 5 of the *Nfe2l2* gene which encodes Nrf2 (Nrf2^flox^; Stock No. 025433) were purchased from Jackson Laboratory (Bar Harbor, Maine). A mouse model with the conditional human heme oxygenase 1 (*HMOX1*) overexpression cassette knocked in the *Rosa26* loci, known as the R26-(CAG-LNL-HMOX)1 mouse, was newly created by Shanghai Biomodel Organism Science & Technology Development Co., Ltd (Shanghai, China). The targeting vector and targeting strategy are illustrated in **Supplementary Figure S1**. This mouse model allows tissue-specific overexpression of HMOX1 when the loxp-flanked expression blocker sequence (“LNL”) is removed by a transgenic Cre that is expressed in the tissue, in which HMOX1 overexpression is controlled by the CAG promoter.^31^ We confirmed cardiac overexpression of the HMOX1 protein in mice harboring both the HMOX1 and the Myh6-Cre transgenes (**Supplementary Figure S2**). Genotypes of mice were determined with PCRs using toe or tail DNA and specific primers (**Supplementary Table S1**).

The animal care and use protocols (12-12-12-15D, 01-01-16-19D) for this study were approved by the Institutional Animal Care and Committee of the University of South Dakota and followed the NIH guide for the care and use of laboratory animals.

Mice were either used for Kaplan-Meier survival analyses or euthanized at 2 or 3 weeks of age for tissue sampling. Unless specified otherwise, mouse ventricular myocardium was stored in RNA-Later for subsequent RNA extraction, snap-frozen in liquid nitrogen and stored in −80°C for subsequent protein analyses, or perfusion-fixed in situ for histopathological assessment.

### Evan’s blue dye (EBD) uptake assay

Detection of CM necrosis in mouse hearts was performed as reported.^8^ In brief, at 3 weeks of age when the homozygous Cops8^CKO^ mice begin to show massive CM necrosis,^7^ mice were injected with EBD (100 mg/kg, i.p.). Eighteen hours after injection, the mice were anesthetized via isoflurane inhalation; in situ retrograded perfusion-fixation via the abdominal aorta was carried out sequentially with 0.9% normal saline and 3.8% paraformaldehyde dissolved in phosphate-buffered saline (PBS). The atria were trimmed, and the fixed ventricles were processed for OCT embedding and subjected to cryosectioning. A series of 7-μm cryosections were collected from the base to the apex of the ventricles. One in every 50 sections was stained for F-actin with Alexa-488 conjugated phalloidin to identify CMs and subjected to imaging with a confocal microscope (Olympus Fluoview 500). The images of each ventricular tissue ring were reconstructed by overlapping images from individual fields and used for quantification of EBD-positive area (red fluorescence) and total F-actin positive area (green fluorescence).

### Necrostatin-1 (Nec-1) treatment

At 2 weeks of age, Cops8^CKO^ mice were continuously administered Nec-1 (BML-AP309, Enzo Life Science; 1.56 mg/kg/day) or vehicle (10% DMSO in PBS) by intraperitoneal implantation of osmotic mini-pumps (Alzet Model 1002, designed for continuous drug delivery for 2 weeks). Two cohorts of mice were included. For CM necrosis analysis using the EBD uptake assay as described above, one cohort of mice was sacrificed 7 days after implantation of the mini-pump. The other cohort was used for Kaplan-Meier survival analysis.

### N-acetyl-L-cysteine (NAC) treatment

At 2 weeks of age, Cops8^CKO^ mice were injected daily for 7 consecutive days with NAC (100 mg/kg/day, i.p.) or vehicle (PBS, pH7.2) before they were subjected to the EBD uptake assay as described above.

### Dihydroethidium (DHE) staining for reactive oxygen species (ROS)

Mouse hearts were perfused *in situ* and excised in PBS, embedded in OCT and rapidly frozen. Serial cryosections (10 μm thick) were mounted onto glass slides. The slides were air-dried and incubated with 2.5 μM DHE (12013, Cayman Chemical, USA) in PBS at 37°C for 30 min. DHE produces a red fluorescence when oxidized to ethidium bromide by the superoxide anion.^32^ The slides were then examined and imaged with a confocal microscope (Olympus Fluoview 500) using a 20X objective. Three mice per genotype, 5 representative tissue sections per heart, and 2 micrographs randomly collected from each section were analyzed. The average density of fluorescence derived from DHE in each confocal micrograph was used as the indicator of ROS content.

### Western blot analyses

Total proteins were extracted from frozen myocardium. Protein concentration was measured using the BCA assay. Proteins fractionated via SDS-PAGE were electro-transferred onto PVDF membrane, immuno-probed for specific proteins using primary and horseradish peroxidase-conjugated secondary antibodies, detected with the enhanced chemiluminescence (ECL) method (RPN2235, Fisher Scientific, USA) as previously reported.^33^ The stain-free total protein imaging technology was used to collect in-lane loading controls for experiments, when appropriate.^34^ The antibodies used include anti-COPS8 antibody (rabbit, BML-PW8290-0100, Enzo Life Science Inc., USA), anti-RIPK1 antibody (mouse, ab72139, Abcam, USA), anti-RIPK3 antibody (rabbit, 14401s, Cell Signaling Technology, Inc., USA), anti-MLKL antibody (rabbit, ab194699, Abcam, USA), anti-Tubulin antibody (mouse, 10806, Sigma-Aldrich, USA), anti-DNP antibody (rabbit, 71-3500, Invitrogen, USA), anti-α-Actinin antibody (mouse, A7811, Sigma-Aldrich, USA), anti-Cullin 3 antibody (rabbit, NB100-58788, Novus, USA), anti-Nrf2 antibody (rabbit, sc-722, Santa Cruz Biotechnology, Inc., USA), anti-phospho-Nrf2 (Ser40) antibody (rabbit, PA5-67520, Invitrogen, USA), anti-KEAP1 antibody (rabbit, 10503-2-AP, Proteintech Group, Inc., USA), and anti-caspase 8 antibody (rabbit, 4790s, Cell Signaling Technology, Inc., USA). BioRad VersaDoc 3000 or ChemiDoc MP and associated QuantityOne or ImageLab softwares (BioRad, Hercules, California, USA) were used for imaging and analyzing chemiluminescence and gel fluorescence.

### Co-immunoprecipitation (Co-IP) assays

The co-immunoprecipitation was performed as previously described.^35^ In brief, protein A/G PLUS-Agarose beads (sc-2003, Santa Cruz Biotechnology Inc., USA) were washed with a buffer (WGB buffer) containing 0.05M Hepes, 0.15M NaCl, and 1% Triton X-100 (pH 7.6) 3 times before being incubated with either anti-RIPK1 antibodies or control IgG for 2 hours at room temperature. The beads were then incubated at 4□ overnight with the crude proteins extracted from ventricular myocardium in the radioimmunoprecipitation assay (RIPA) buffer. The beads were then spun down, separated from supernatant, and further washed 3 times (5 min per wash) with the WGB buffer to remove unbound proteins; proteins bound on the beads were then eluted with SDS loading buffer (50 mM Tris-HCl at pH 6.8, 2% SDS, and 10% glycerol) and then boiled for 5 min. The eluted proteins were subjected to SDS-PAGE and western blot analyses for RIPK1 and RIPK3 with the western blot protocol as described above.

### Protein carbonyl assays

Protein carbonyl assays used the Oxidized Protein Western Blot Detection Kit (ab178020; Abcam, USA) and were performed as we previously described.^36^ Briefly, ventricular myocardium was homogenized in RIPA buffer. After centrifugation, the supernatant was collected and supplemented with DTT (50 mM, final concentration). Protein samples were then mixed with the same volume of 12% SDS and incubated with an equal volume of the 1× 2,4-dinitrophenylhydrazine (DNPH) derivatization solution at room temperature for 15 min before reaction termination by addition of the neutralization solution. The carbonyl groups in the protein side chains are derivatized to 2,4-dinitrophenylhydrazone (DNP-hydrazone). The DNP-derivatized proteins were then subjected to SDS-PAGE and western blot analysis or loaded directly onto a PVDF membrane via a vacuum-assisted device and detected using dot blotting with an anti-DNP antibody.

### Caspase 8 activity assays

The activities of caspase 8 in myocardial crude protein extracts were measured using the Caspase-8 Colorimetric Assay Kit (K113, BioVision, Inc., USA).

### Statistical analyses

The presentation of quantitative data and the methods for statistical analyses are described in the legend of each figure.

## Results

### Key proteins of the necroptotic pathway are increased in Cops8^CKO^ mouse hearts

We have previously observed massive CM necrosis in mice with Cops8^CKO^ initiated at either the perinatal or adult stage.^7, 9^ To explore the mechanism governing the CM necrosis in Cops8 deficient hearts, we examined the potential involvement of the RIPK1-RIPK3 pathway. Western blot analyses revealed marked increases in myocardial protein levels of RIPK1, RIPK3, and MLKL in mice with perinatal Cops8^CKO^ compared with littermate control mice (**Figure 1A, 1B**). Co-immunoprecipitation of RIPK1 detected increased association of RIPK3 with RIPK1 in Cops8^CKO^ hearts compared with littermate controls (**Figure 1C, 1D**). Increased RIPK1-RIPK3 interaction is a key step in the activation of the necroptotic pathway by death receptor engagement;^37–39^ hence, these data suggest that the RIPK1-RIPK3 pathway is likely activated in Cops8 deficient hearts.

**Figure 1.**
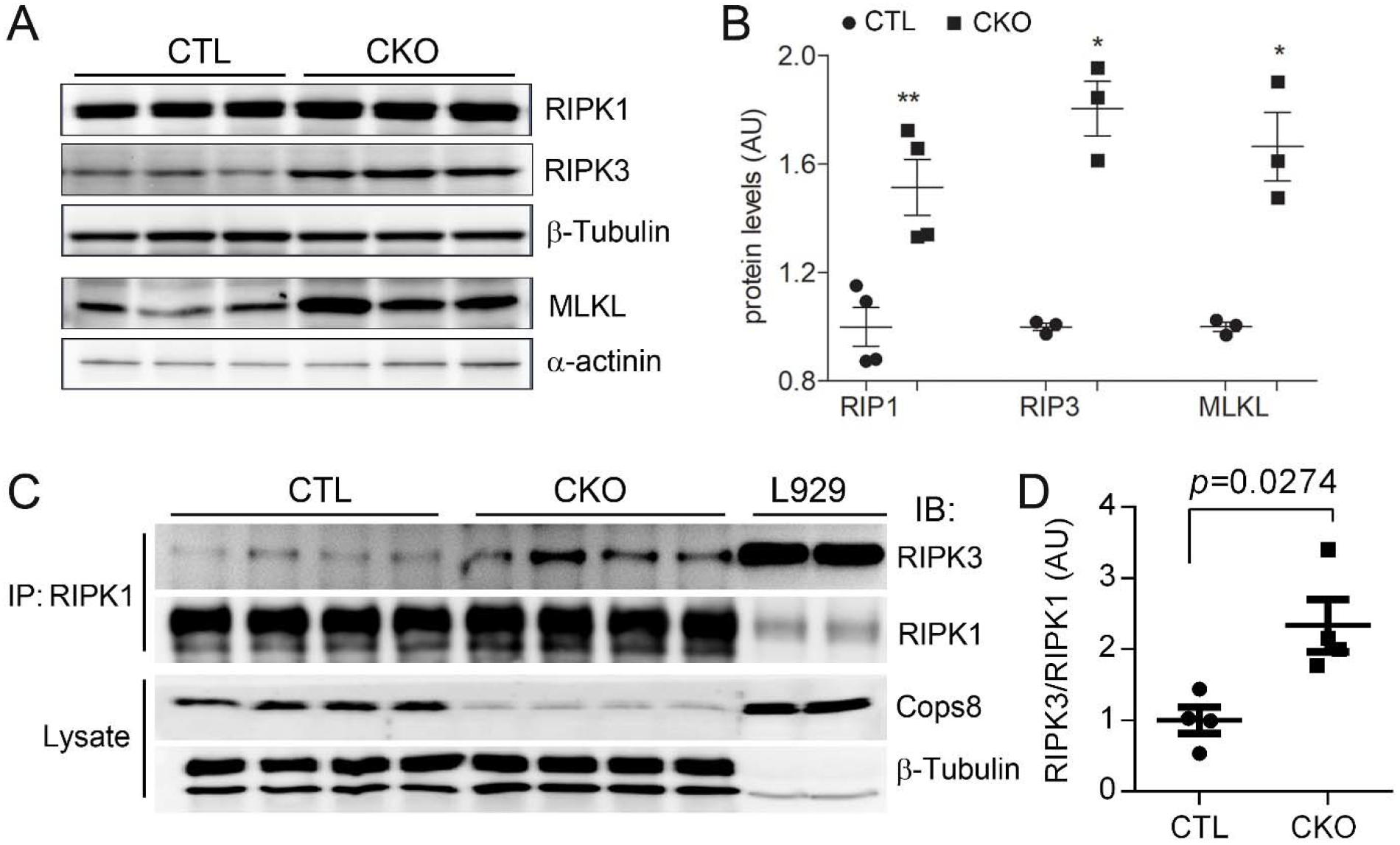
Increases in myocardial RIPK1, RIPK3 and MLKL and in RIPK1-bound RIPK3 proteins in Cops8^CKO^ mice. **A** and **B**, Representative images (A) and pooled densitometry data (B) of western blot analysis for the indicated proteins in the myocardial extracts of 3-week-old Cops8^CKO^ (CKO) and littermate control (CTL) mice. β-Tubulin and α-actinin were probed as loading controls for the proteins shown above. Mean with SEM, \**p*<.05, **p<0.01 vs. CTL; **C**, Western blot (IB) analyses for RIPK1 and RIPK3 in the RIPK1 immuno-precipitates (IP) from the protein lysate of ventricular myocardium from 3-week-old CTL and Cops8^CKO^ mice. One mouse/lane. L929 cell lysates were used as positive controls. **D**, RIPK1/RIPK1 ratios in the RIPK1 IP. The density of RIPK3 and RIPK1 bands for individual samples shown in panel C was used for the calculation of RIPK3 to RIPK1 ratios, the mean of the ratios of the CTL group is defined as 1 arbitrary unit (AU). The *p* values shown in this figure are derived from two-side unpaired *t*-test with Welch’s correction.

### Suppression of CM necrosis and delay of premature death by RIPK1 inhibition in Cops8^CKO^ mice

To determine whether RIPK1 kinase activity is required for CM necrosis in Cops8^CKO^ hearts, we tested the impact of necrostatin-1 (Nec-1), a RIPK1 kinase-specific inhibitor.^40^ Since CM necrosis is detectable at 3 weeks of age, but not at 2 weeks, the administration of Nec-1 or vehicle control via intraperitoneal implantation of osmotic mini-pumps was initiated in Cops8^CKO^ mice at 2 weeks of age. CM necrosis was assessed with the in vivo EBD uptake assay in the heart harvested 7 days after mini-pump implantation. EBD positive CMs were not detectable in mice with control genotypes (Myh6-Cre^TG^, Cops8^FL/FL^, and Cops8^+/+^; data not shown) but were readily detectable in homozygous Cops8^CKO^ mice treated with vehicle control. Strikingly, the EBD positivity in Cops8^CKO^ mouse hearts was nearly abolished completely by the Nec-1 treatment (**Figure 2A, 2B**, *p*<0.0001), indicating that RIPK1 kinase activity is required for Cops8 deficiency to induce CM necrosis in mice. Moreover, Kaplan-Meier survival analyses revealed that Nec-1 treatment significantly delayed the premature death observed in Cops8^CKO^ mice (*p*=0.0072, **Figure 2C**). Taken together, these findings provide compelling evidence that induction of CM necrosis by Cops8 deficiency requires RIPK1 kinase activity and the CM necroptosis is principally responsible for the premature death of Cops8^CKO^ mice.

**Figure 2.**
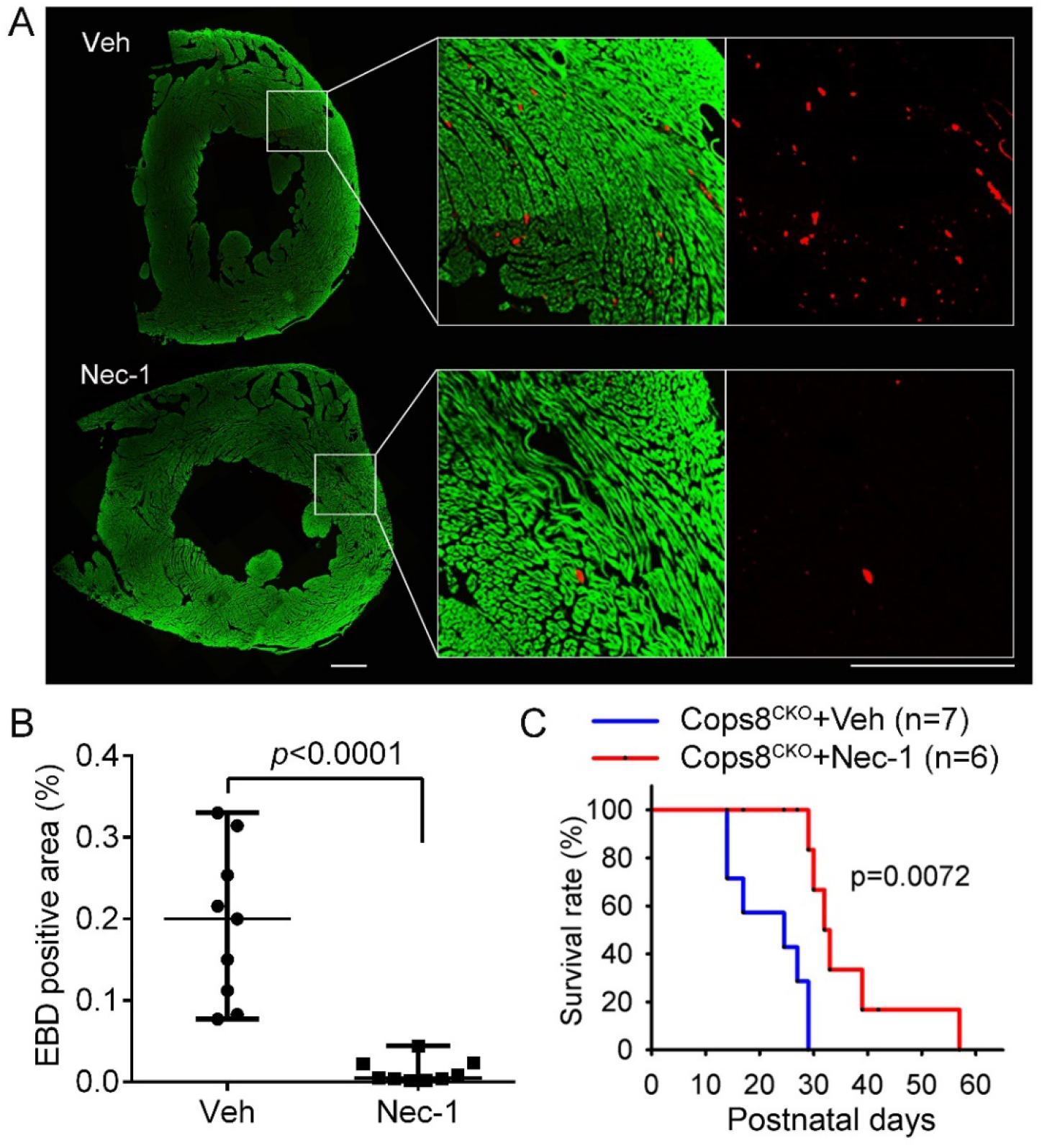
Necrostatin-1 (Nec-1) treatment markedly reduces CM necrosis and delays premature death of Cops8^CKO^ mice. Cohorts of Cops8^CKO^ mice at 2 weeks of age were treated with necrostatin-1 (Nec-1, 1.56 mg/kg/day) or vehicle (Veh) via intraperitoneal osmatic mini-pumps for 1 week (**A**, **B**) or continued for >2 weeks for the Kaplan-Meier survival analysis (**C**). **A** and **B**, At day 6, after min-pump implantation, mice were treated with one dose of Evan’s blue dye (EBD; 100 mg/kg, i.p.) 18 hours before they were anesthetized and perfusion-fixed in situ. Cryosections from the fixed heart were stained with Alexa488-conjugated phalloidin to identify CMs (green) and subjected to fluorescence confocal imaging analyses. The images of each ventricular tissue ring were reconstructed and used for quantification of EBD-positive area (red) and total green area. Panel **A** shows representative reconstructed images from a pair of Cops8^CKO^ hearts treated with Veh or Nec-1; scale bar=0.5 mm. Individual percent values of average EBD positive area in the 3 representative sections/mouse from 3 mice of each group are plotted in panel **B**, superimposed by median with range; Mann Whitney test. **C**, Kaplan-Meier survival curve of Cops8^CKO^ mice treated with Veh or Nec-1. Nec-1 treatment significantly increased lifespan of Cops8^CKO^ mice compared with the vehicle-treated group (median lifespan: 32.5 vs. 27 days); Log-rank Test.

### Requirement of RIPK3 for CM necrosis in Cops8^CKO^ mice

To test the role of RIPK3 in the CM necrosis of Cops8^CKO^ mice, RIPK3 germline knockout (RIPK3^−/−^) mice were cross-bred with Cops8^CKO^ mice and the resultant Cops8^CKO^::RIPK3^+/+^ and Cops8^CKO^::RIPK3^+/−^ littermate mice were subjected to EBD CM necrosis assessment at 3 weeks of age as well as Kaplan-Meier survival analysis. The prevalence of EBD-positive CMs in Cops8^CKO^::RIPK3^+/−^ mice was significantly lower than that of littermate Cops8^CKO^::RIPK3^+/+^ mice (*p*=0.0007; **Figure 3A, 3B**;); also, the lifespan of the former was significantly longer than that of the latter (*p*<0.0001; **Figure 3C**). These analyses show that RIPK3 haploinsufficiency is capable of markedly suppressing CM necrosis and delaying premature death in Cops8^CKO^ mice, providing compelling evidence that RIPK3 is required for CM necrosis in Cops8^CKO^ mice. The findings described so far also demonstrate that CM necrosis induced by Cops8 deficiency belongs to necroptosis and is mediated primarily by the RIPK1-RIPK3 pathway.

**Figure 3.**
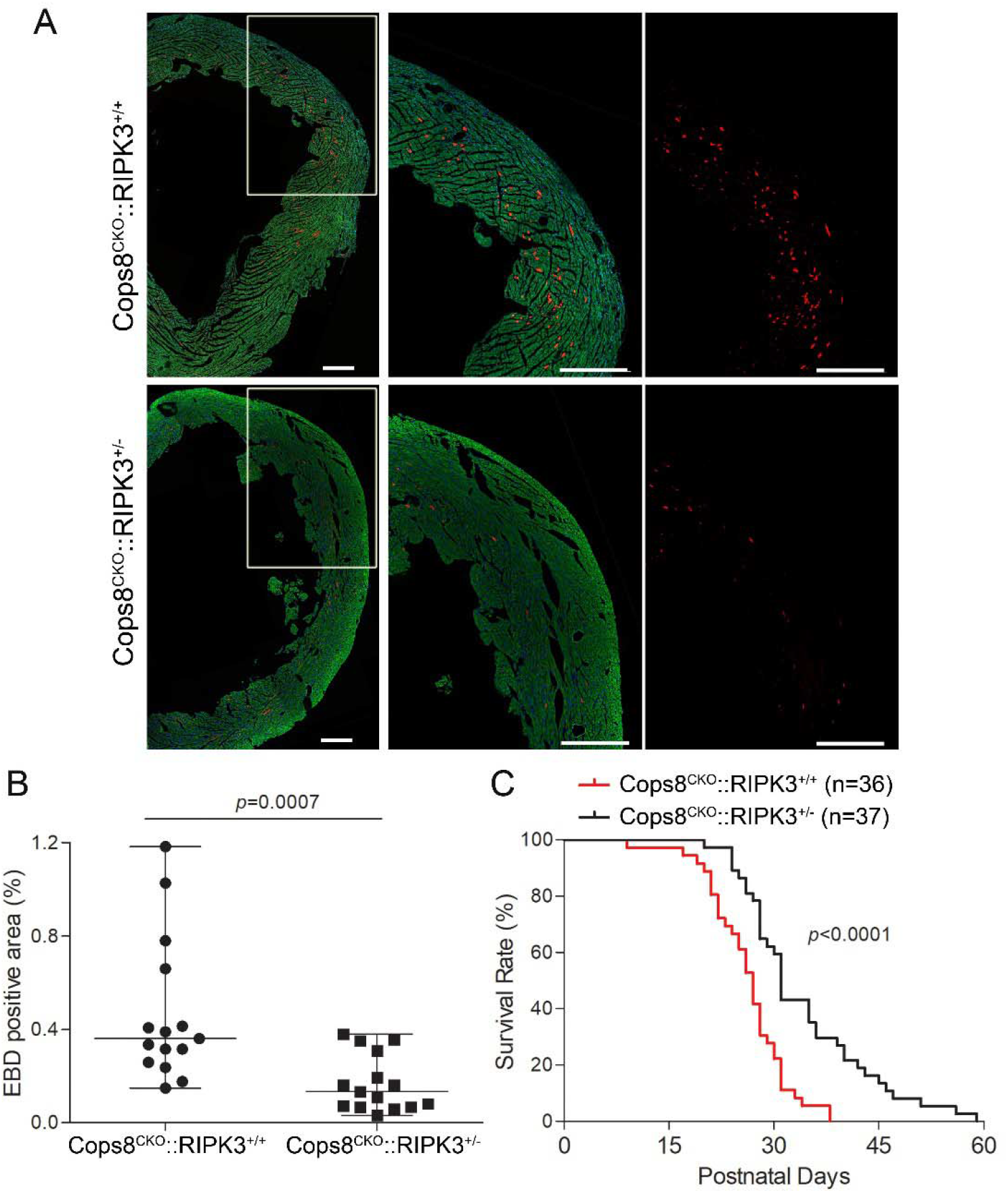
RIP3 haploinsufficiency significantly reduces CM necrosis and delays premature death of Cops8^CKO^ mice. **A**, Representative confocal micrographs of EBD assays. Littermate mice of the indicated genotypes at 3 weeks of age were subjected to the EBD assays in the same way as described in Figure 2. EBD positive cells display autofluorescence (red) and F-actin was stained using Alexa-488-conjugated phalloidin (green). Shown are representative composed images for the entire cross-section of the left ventricle or a higher magnification view of the marked portion of the composed image (**A**). Scale bar=500μm. **B**, dot plot to show the individual percent values of EBD positive area in the 5 representative sections/mouse of 3 mice of each group. Median with range is superimposed. Mann Whitney test. **C**. Kaplan-Meier survival curve. RIPK3 haploinsufficiency (RIPK3^+/−^) delayed premature death of Cops8^CKO^ mice. Log-Rank Test.

### CM necroptosis in Cops8^CKO^ mice is independent of mitochondrial permeability transition (MPT)

By definition, necroptosis and MPT-driven necrosis are two different types of regulated necrosis;^41^ however, it was previously reported that Nec-1 failed to show additional protection against myocardial I/R injury in Cyclophilin D knockout (*Ppif^−/−^*) mice,^42^ inferring that MPT and RIPK1 might be involved in the same regulatory pathway. More recently, MPT was shown as a major player in the RIPK3-CaMKII-MPT pathway for the induction of myocardial necroptosis by I/R and doxorubicin.^23^ Hence, we determined whether MPT-driven necrosis contributes to CM necrosis in Cops8^CKO^ mice by testing whether ablation of the *Ppif* gene would mitigate the CM necrosis and mouse premature death induced by Cops8^CKO^. As presented in **Figure 4**, neither heterozygous nor homozygous knockout of the *Ppif* gene delayed the mouse premature death; on the contrary, homozygous *Ppif* knockout moderately increased CM necrosis (*p*=0.010) and accelerated mouse premature death (*p*=0.007), indicating that MPT is not a mediator for CM necrosis in Cops8^CKO^ mice.

**Figure 4.**
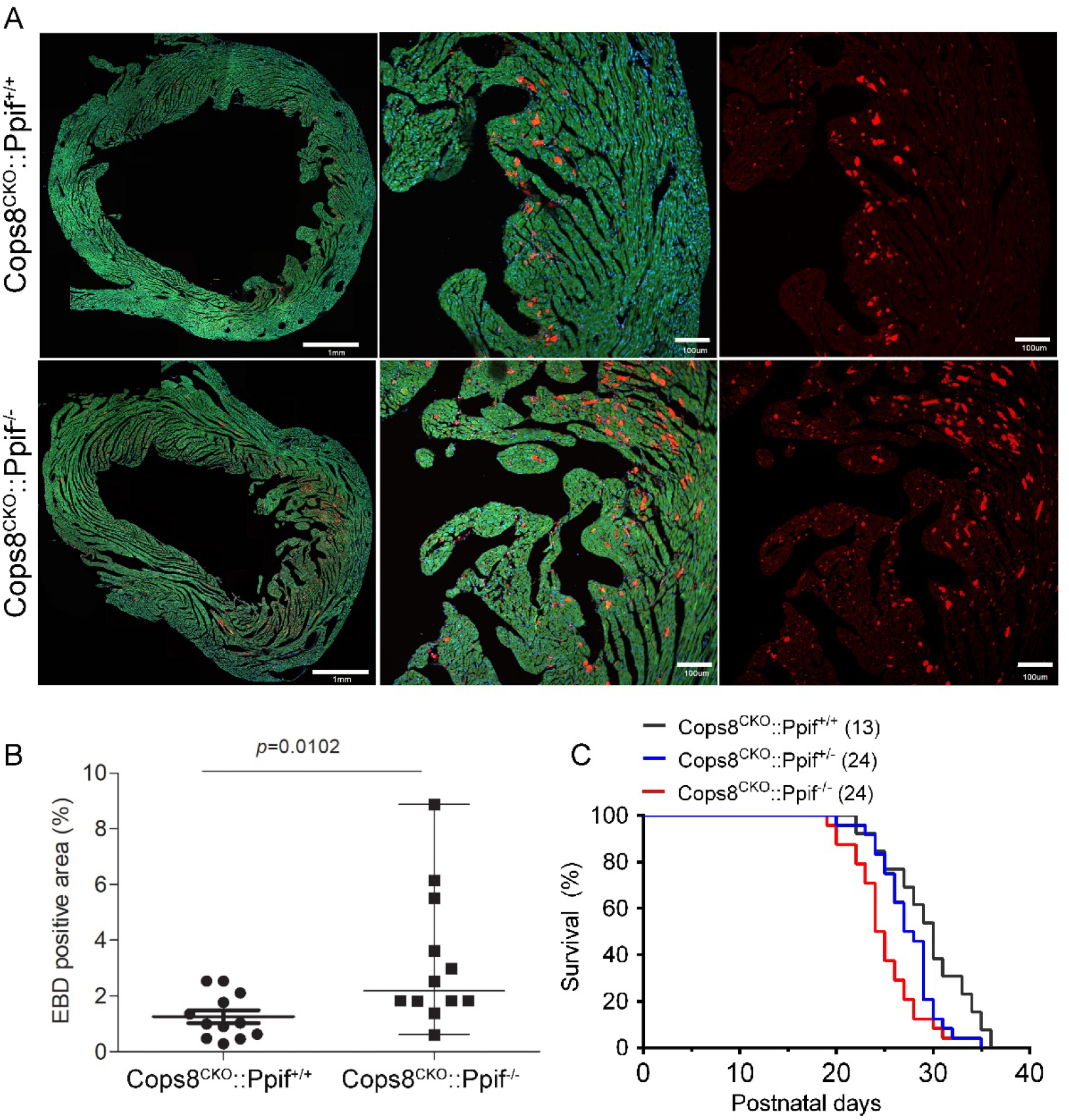
Cyclophilin D knockout exacerbates CM necrosis and premature death of Cosp8^CKO^ mice. The *Ppif* gene encodes Cyclophilin D. Cops8^CKO^ was induced in the *Ppif* wild type as well as hemizygous and homozygous null background through cross-breeding. **A** and **B**, in vivo EBD uptake assays were performed on mice at 3 weeks of age as described in Figure 2. Shown are representative confocal micrographs of myocardial sections from mice of the indicated genotypes (A) and scatter dot plot of the individual percent values of average EBD positive area in the 3 representative sections per mouse and 4 mice per group, superimposed by median with range (B). Mann Whitney test. **C**, Kaplan-Meier survival curves of littermate mice with the indicated genotypes. *p*=0.007, Cops8^CKO^::Ppif^−/−^ vs. Cops8^CKO^::Ppif^+/+^, Log-rank test.

### Cops8 deficiency increases myocardial oxidative stress but ROS scavenging fails to suppress CM necroptosis in Cops8^CKO^ mice

The level of superoxide anion (O_2_) in myocardial sections was probed with DHE incubation followed by fluorescence confocal microscopy. Upon exposure to superoxide anion, DHE is converted to 2-hydroxyethidium, which then intercalates into nuclear DNA and exhibits red fluorescence.^32^ The red fluorescence intensity of the DHE-probed myocardial sections from homozygous Cops8^CKO^ mice was remarkably greater than that from either Cops8^FL/+^::Myh6-cre^TG^ (heterozygous Cops8^CKO^) or Cops8^FL/FL^ control mice (**Figure 5A, 5B**), indicating that Cops8 deficiency increases myocardial superoxide levels. Myocardial reactive oxygen species (ROS) were also assessed via immunoblotting for DNPH-derivatized protein carbonyls. Immuno-probing of DNP in protein dot blots revealed that myocardial protein carbonyls were substantially higher in the homozygous Cops8^CKO^ mice compared with heterozygous Cops8^CKO^, Cops8^FL/FL^, or Myh6-Cre^TG^ mice (**Figure 5C, 5D**). Western blot analyses further showed that the increased carbonyls were mainly on proteins of a molecular weight ranging from 25 to 37 kDa (**Figure 5E**). These findings indicate that Cops8 deficiency in CMs increases myocardial oxidative stress.

**Figure 5.**
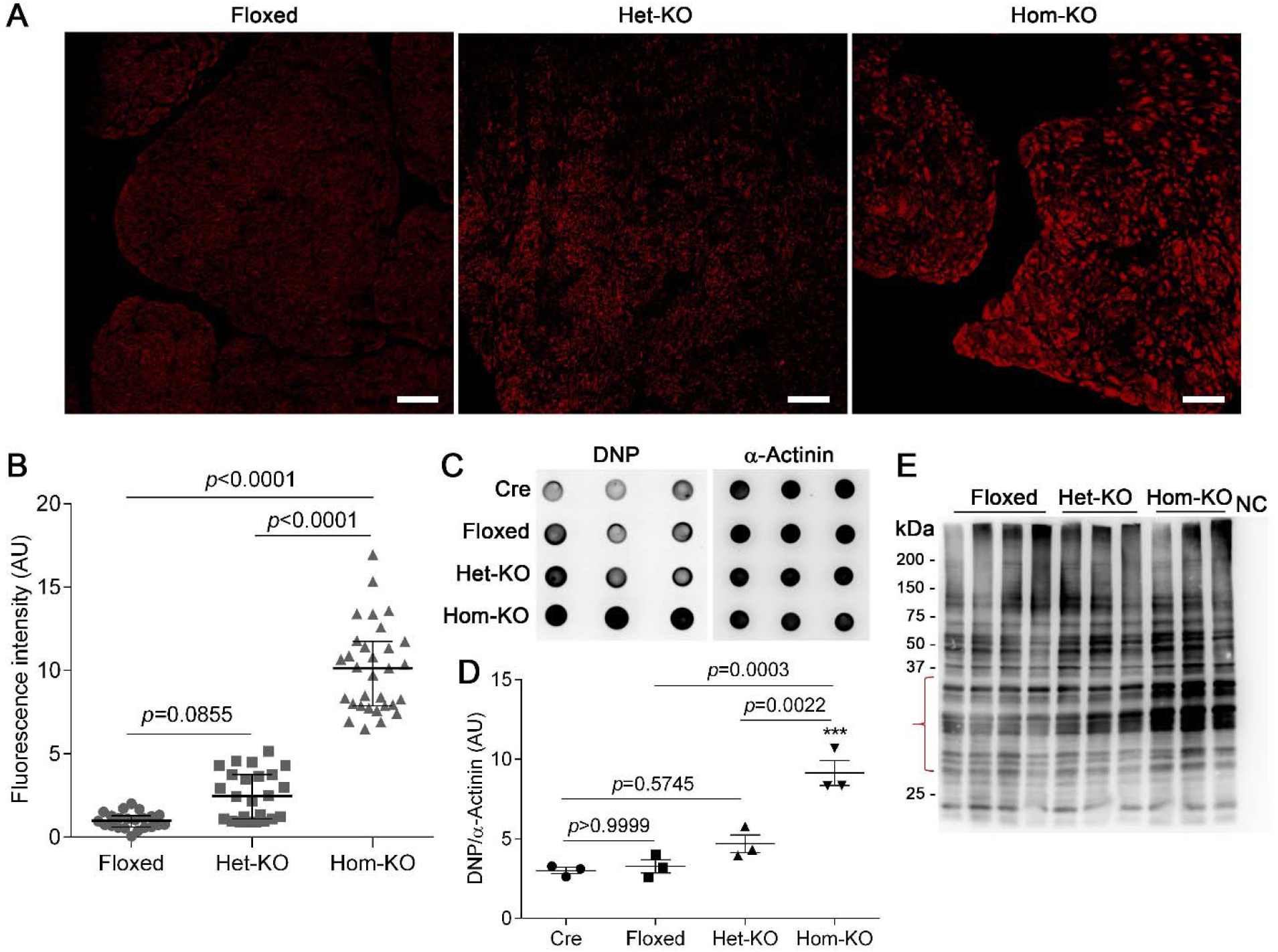
Changes of myocardial reactive oxygen species (ROS) in Cops8^CKO^ mice. **A** and **B**, Detection of ROS in myocardial sections by dihydroethidium (DHE) staining (red). Four to six representative sections per mouse and 5 mice per genotype were analyzed. Panel A shows representative confocal fluorescence images of DHE stained myocardial sections from homozygous Cops8^FL/FL^, heterozygous Cops8^CKO^ (Het-KO), and homozygous Cops8^CKO^ (Hom-KO) mice. Scale bar=100μm. Panel B presents a scatter dot plot of individual average fluorescent intensity values for each sections, superimposed by median with interquartile range. The non-parametric Kruskal-Walls test followed by Dunn’s multiple comparison tests were used. **C** and **D,** Dot blot analyses for DNP-derivatized protein carbonyls. Equal amounts of proteins were subject to DNP-derivatization and equal proportions of the DNP-derivatized preparation were used for dot blot and subsequent immunoprobing for DNP. α-Actinin was probed as a loading control. Shown are representative images (**C**) and pooled densitometry data (**D**). Mean±SEM are superimposed. Cre, Myh6-Cre^TG^ only; \*\*\**p* =0.0002 vs. Cre, one way ANOVA followed by Bonferroni’s multiple comparisons test. **E**, Representative image of western blot analysis of DNP-derivatized protein carbonyls. The opening curly brace demarcates the protein molecular weight range where carbonyls were increased most in the Hom-KO group. NC, negative control where equal amount of myocardial proteins that were not subject to DNP derivatization was loaded.

Increased oxidative stress is considered a main factor for causing necroptosis. Since ROS was remarkably increased in Cops8^CKO^ hearts, we sought to determine its contribution to the necroptosis by examining the impact of treatment with N-acetyl-cysteine (NAC), a widely used free radical scavenger, on the CM necrosis. Unexpectedly, NAC treatment failed to reduce EBD positivity in Cops8^CKO^ hearts; on the contrary, it moderately increased CM necrosis (*p*=0.017; **Figure 6A, B**). Heme oxygenase 1 (HMOX1) is an antioxidant. We next further tested whether a genetic method to increase anti-oxidative capacity in CMs would be effective in modulating the Cops8^CKO^ phenotype by transgenic overexpression of HMOX1 in CMs. Kaplan-Meier survival analysis showed that cardiomyocyte-restricted overexpression of HMOX1 did not delay the premature death of Cops8^CKO^ mice. On the contrary, the HMOX1 overexpressed Cops8^CKO^ mice tended to show a shorter lifespan (*p*=0.044; **Figure 6C**). Taken together, these data indicate that increasing reductive capacity via either pharmacological or genetic means tend to exacerbate cardiac pathology in Cops8^CKO^ mice.

**Figure 6.**
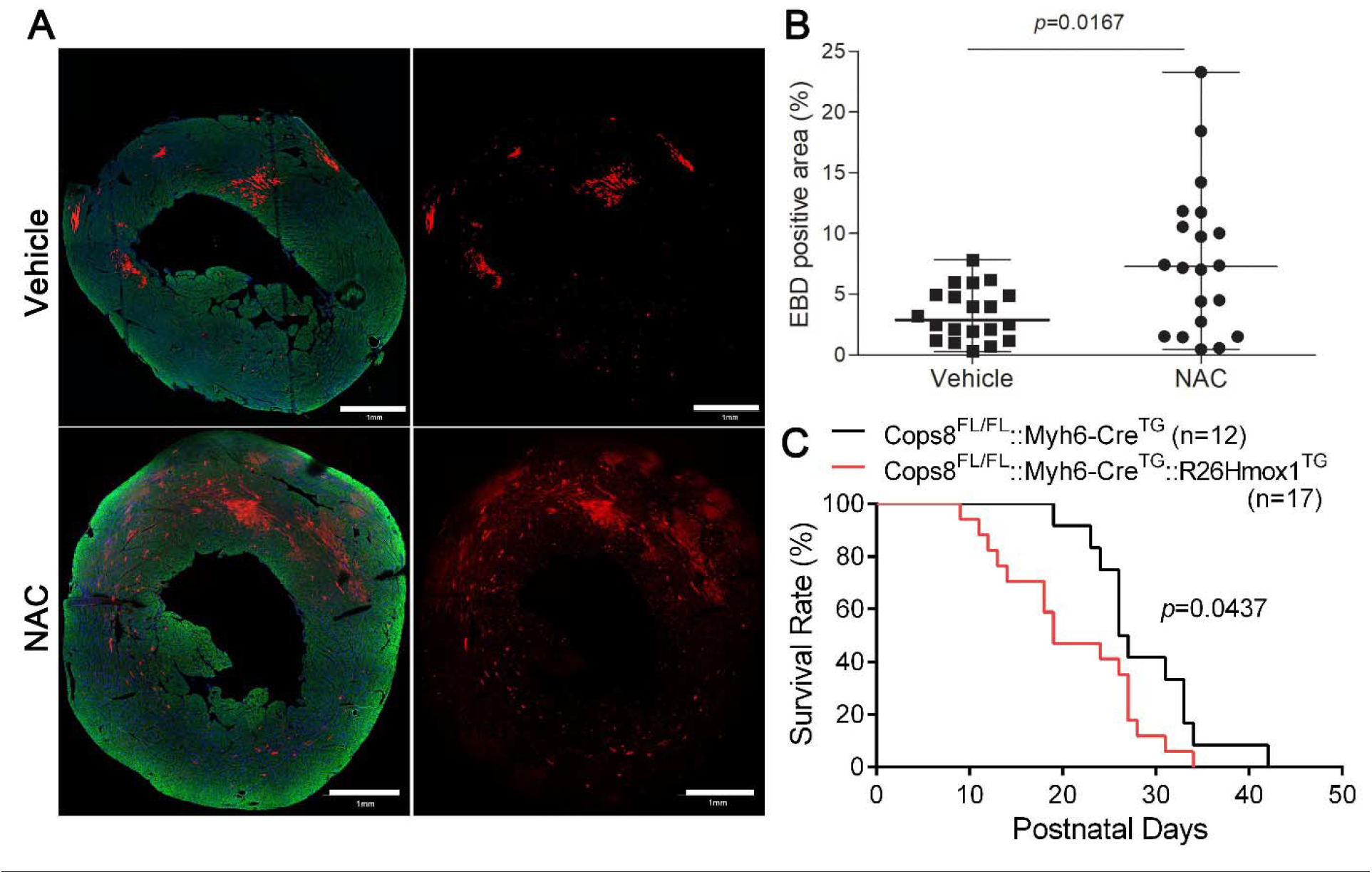
NAC treatment and Hmox1 overexpression exacerbates CM necrosis and premature death in Cops8^CKO^ mice. **A** and **B**, Representative composed confocal images (A) and pooled quantitative data (B) from the EBD uptake assays for LV myocardium of Cops8^CKO^ mice treated with NAC or vehicle control. Seven consecutive daily intraperitoneal injections of NAC (100 mg/kg/day) or vehicle were initiated at 14 days of age. EBD assays were performed at 21 days of age as described in Figure 2. EBD positive CMs emit auto-fluorescence (red); Alexa fluor-488-conjugated phalloidin was used to stain F-actin and thereby identify cardiomyocytes (green). In the dot plot (B), individual percent values of average EBD positive area are shown. Four representative sections/mouse and 5 mice/group were included. Median with range, *p*=0.0167, Mann Whitney test. **C**, Kaplan-Meier survival curve of mice of the indicated genotypes. Both males and females (roughly 1:1 ratio) were included. Log-rank test.

### Impaired caspase 8 activation and upregulated BCL2 in Cops8^CKO^ hearts

Since necroptosis was originally observed in TNFα-treated cells whose caspase 8 is defective or suppressed, we sought to examine myocardial expression and activity of caspase 8 in Cops8^CKO^ mice. Both the cleaved/activated form of caspase 8 and the activities of caspase 8 were markedly lower but the abundance of the full-length caspase 8 was discernibly greater in the Cops8^CKO^ hearts compared with littermate controls at 3 weeks of age (**Figure 7A ~ 7C**), which indicates that cardiac Cops8 deficiency suppresses caspase 8 activation; thereby, Cops8 deficiency suppresses the activation of the extrinsic apoptotic pathway. As we reported before, myocardial levels of BCL2, a key inhibitor of the mitochondrial apoptotic pathway, were significantly increased in 3-week-old homozygous Cops8^CKO^ mice, compared with littermate control mice with heterozygous Cops8^CKO^ and Cops8^FL/FL^ littermates (*p*=0.0102, 0.0003; **Figure 7D**). Myocardial BCL2 mRNA levels were also greater in homozygous Cops8^CKO^ mice than littermate controls at both 2 and 3 weeks of age (**Figure 7E**). Taken together, these data support that Cops8 deficiency suppresses apoptotic pathways.

**Figure 7.**
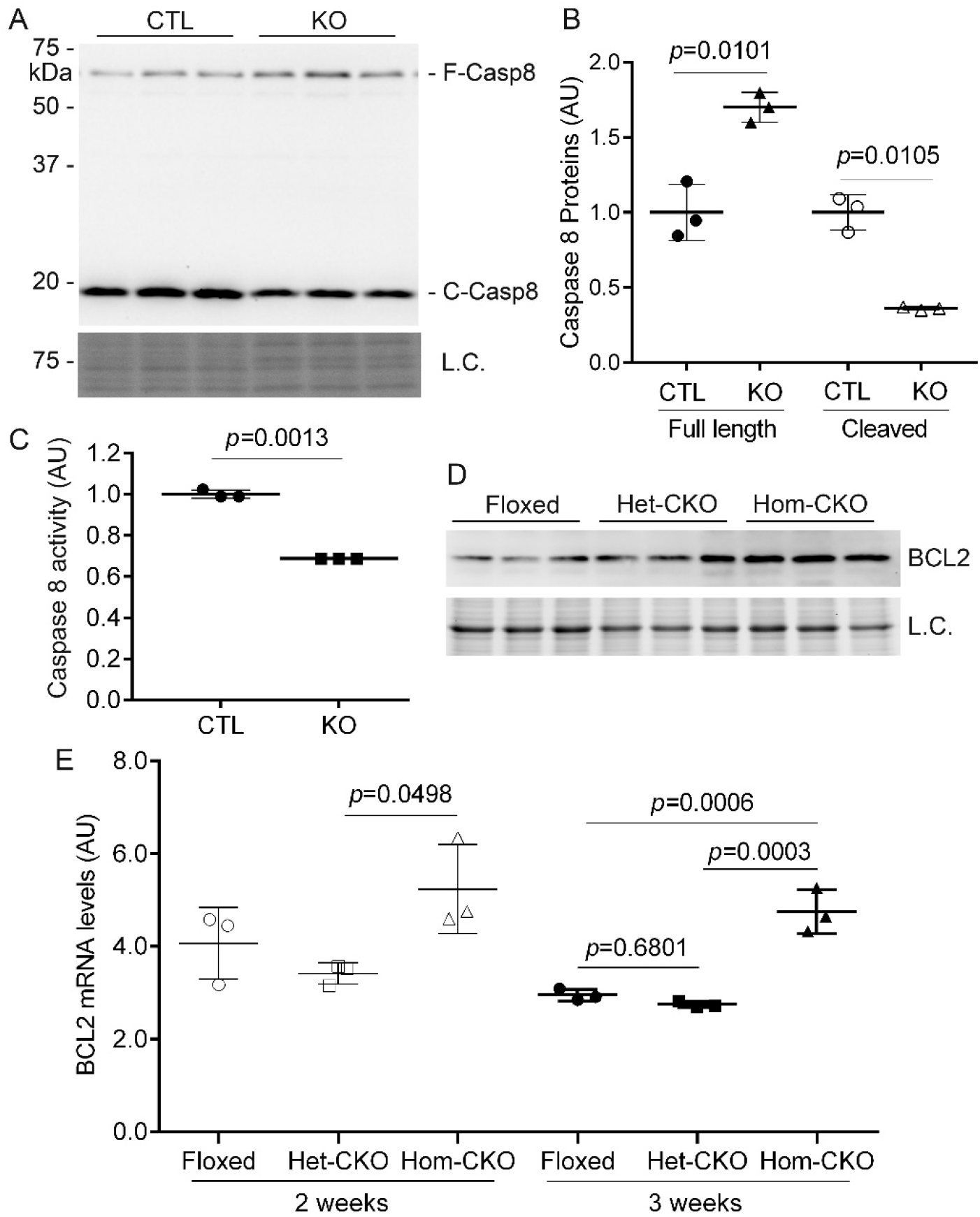
Changes in both protein expression and activities of caspases 8 as well as BCL2 protein and mRNA levels in Cops8^CKO^ mouse hearts. **A** and **B**, Representative images (A) and scatter dot plots of pooled densitometry data (B) of western blot analyses for caspase 8 (Casp8). L.C., Loading control which is a portion of the image from stain-free in-gel imaging of total proteins that was used to normalize caspase 8 western blot signals. F-, full length; C-, cleaved form. **C**, Changes in myocardial caspase 8 activities in Cops8^CKO^ mice at 3 weeks. CTL, littermate control; KO, homozygous Cops8^CKO^. **D**, Representative images of western blot analyses for myocardial BCL2 in homozygous Cops8^FL/FL^ (Floxed), heterozygous Cops8^CKO^ (Het-CKO), and homozygous Cops8^CKO^ (Hom-CKO) mice at 3 weeks of age. **E**, Changes in myocardial BCL2 mRNA levels in mice at 2 and 3 weeks of age. Each scatter dot plot is superimposed by mean±SD; each dot represents a mouse; *p* values are derived from unpaired t-tests with Welch’s correction (B, C) or one way ANOVA followed by Tukey’s test (E).

### Contributions of increased Nrf2 to CM necroptosis in Cops8^CKO^ mice

Increased oxidative stress is known to activate the nuclear factor E2-related factor 2 (Nrf2). Indeed, our prior transcriptome analysis has revealed that Nrf2 target genes are markedly upregulated in Cops8^CKO^ hearts.^43^ Here our further work detected that myocardial protein levels of total Nrf2 and Ser40-phosphorylated Nrf2 (pS40-Nrf2) were significantly increased in Cops8^CKO^ mice at 2 and 3 weeks of age, compared with littermate controls (**Figure 8A~8C**). Phosphorylation of Nrf2 by protein kinase C (PKC) at Ser40 is known to promote Nrf2 nuclear translocation and increase its target gene expression;^44^ hence, the increases in pS40-Nrf2 are consistent with increased Nrf2 transactivation in Cops8 deficient hearts as we previously detected via transcriptome profiling.^43^

**Figure 8.**
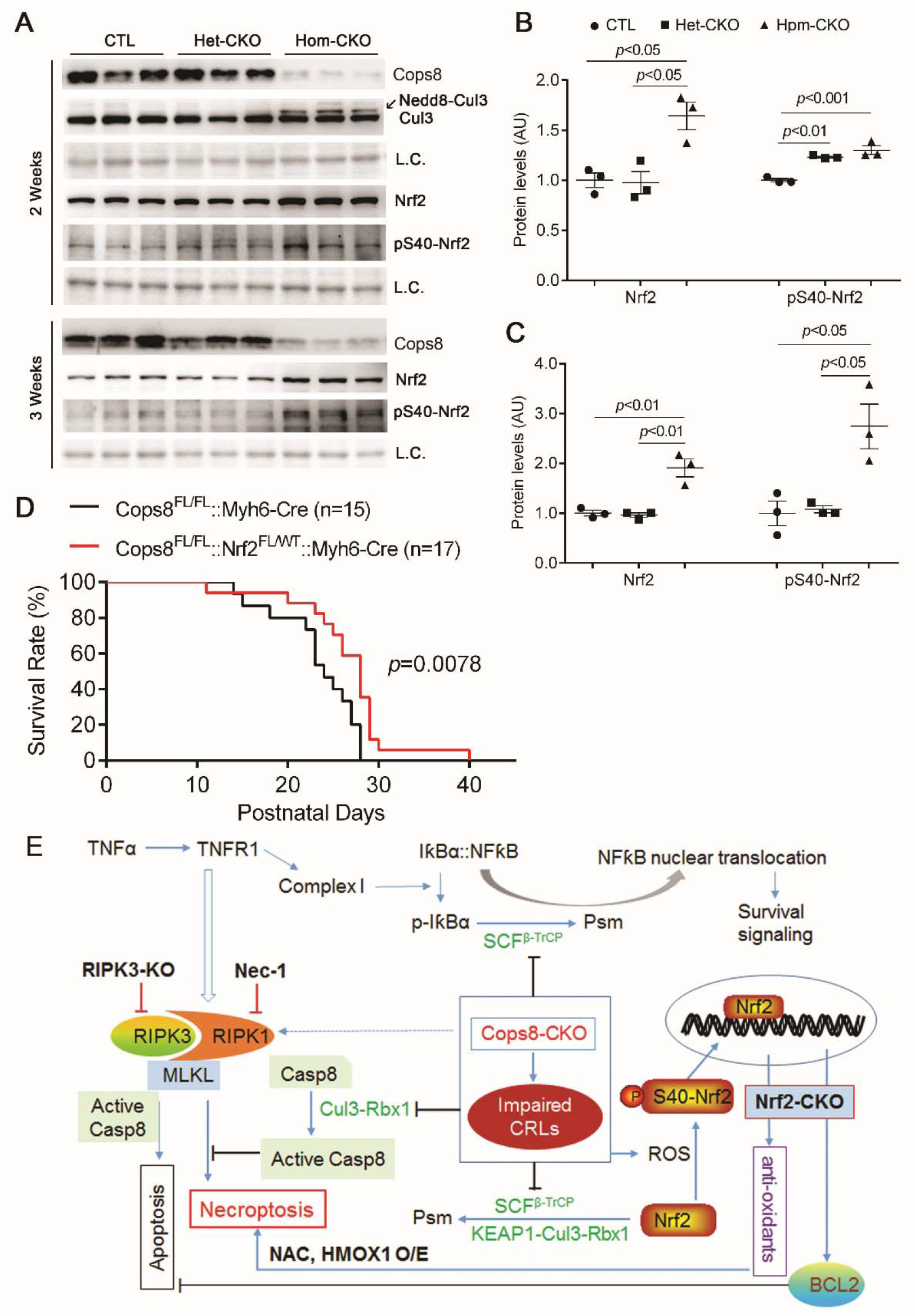
The Nrf2-BCL2 pathway is activated and contributes to CM necroptosis in Cops8^CKO^ mouse hearts. **A** ~ **C**, representative images (A) and the summary of densitometry data (B, C) of western blot analyses for the indicated proteins in the ventricular myocardium of mice of the indicated genotypes at 2 and 3 weeks of age. Here CTL are comprised of Myh6-Cre^TG^ mice. One way ANOVA followed by Tukey’s test. **D**, Kaplan-Meier survival curve of littermate mice of the indicated genotypes. The median lifespan for Cops8^CKO^ mice in the heterozygous Nrf2^CKO^ background (Cops8^FL/FL^::Nrf2^FL/WT^::Myh6-Cre) or in the wild type *Nrf2* background (Cops8^FL/FL^::Myh6-Cre) is respectively 28 or 24 days. Log-rank test. Both male and female (roughly 1:1 ratio) were included in all studies. **E**, A working model for induction of CM necroptosis by Cops8 deficiency, with the main interrogations of this study marked with bold black font. Casp8, caspase 8; dot line denotes a potential link that is not tested yet.

To test the role of increased Nrf2 in the CM necroptosis, we crossbred the Nrf2-floxed allele into Cops8^CKO^ mice and performed Kaplan-Meier survival analysis among the littermates (**Figure 8D**). The lifespan of Cops8^FL/FL^::Nrf2^FL/FL^::Myh6-Cre^TG^ was comparable to, but that of Cops8^FL/FL^::Nrf2^FL/+^::Myh6-Cre^TG^ was significantly longer than, that of Cops8^FL/FL^::Nrf2^+/+^::Myh6-Cre^TG^ mice (*p*=0.0078), indicating that cardiomyocyte-restricted *Nrf2* haploinsufficiency attenuates CM necroptosis induced by CM Cops8 deficiency in mice.

## Discussion

The present study unveils for the first time that CMs deficient of Cops8 die primarily in the form of necroptosis. Mechanistically, by virtue of impairing CRL-mediated ubiquitination, Cops8 deficiency impairs caspase 8 activation and sustains the activation of the Nrf2-BCL2 axis, thereby suppressing both extrinsic and intrinsic apoptotic pathways, which steers the death receptor-mediated signaling towards activation of the RIPK1-RIPK3-mediated necroptotic pathway. Findings of this study also demonstrate that the MPT does not play an important role in CM necroptosis induced by Cops8^CKO^ in mice whereas sustained Nrf2 activation and reductive stress contribute to the induction of CM necrosis and cardiac malfunction by Cops8 deficiency in CMs. These discoveries not only establish the CSN as a crucial factor to suppress CM necroptosis but provide the first demonstration in any organs or systems that, in a UPS and autophagy impairment setting, sustained Nrf2 activation and reductive stress pivot the cardiomyocyte to necroptosis, both of which have highly significant clinical implications.

### Cops8 deficient or CSN inhibited CMs die primarily from necroptosis

Massive CM necrosis occurs in Cops8^CKO^ mice, as evidenced by rapid increases in EBD uptake by CMs in the absence of increased TUNEL positivity, as well as by the ultrastructural features like CM swelling and a broken plasma membrane.^7–9^ Activation of RIPK3 is the centerpiece of necroptotic pathway although RIPK1 is also required in the induction of necroptosis by TNFα at least.^45^ Unlike detection of apoptosis for which a series of relatively simple and specific assays have long been developed, the detection of necroptosis currently requires a combination of rather sophisticate tests to reveal both the necrotic feature (e.g., loss of plasma membrane integrity) and the dependence on RIPK3 activation, according to a recently published guideline.^46^ In the present study, we found that CM necrosis in Cops8^CKO^ mice were associated with increases in myocardial protein levels of RIPK1, RIPK3, MLKL, and RIPK1-bound RIPK3 (**Figure 1**) and were dependent on RIPK1 kinase activity (**Figure 2**) and increased expression of RIPK3 (**Figure 3**), demonstrating unequivocally that the massive CM necrosis observed in Cops8^CKO^ mice belongs to necroptosis. Notably, in contrast to a recently delineated RIPK3-CamKII-MPT pathway to cardiac necroptosis,^23^ MPT does not play a major role in the execution of CM necroptosis in Cops8^CKO^ mice. This is because Cyclophilin D knockout, which is known to inhibit MPT, did not attenuate but rather exacerbated CM necrosis and premature death in Cops8^CKO^ mice (**Figure 4**).

### How does Cops8 deficiency cause CM necroptosis?

The requirement of both RIPK1 and RIPK3 by the CM necrosis observed here suggests that the induction of CM necroptosis by Cops8CKO shares the same pathway taken byTNFR1 activation. The ligation of TNFR1by TNFα can lead to at least 3 possible downstream events: (1) formation of complex 1 where RIPK1 serves as a scaffold in a manner independent of its kinase activity, which provides survival signals via activation of nuclear factor κB (NFκB) and mitogen-activated protein kinases (MAPKs), (2) formation of complex 2a which induces apoptosis via caspase 8 and downstream cascade, and (3) formation of complex 2b (i.e., the RIPK1-RIPK3-MLKL) and thereby induction of necroptosis when caspase 8 is defective or inhibited.^11^ The kinase activity of RIPK1 is required for RIPK1 to induce cell death in complex 2. UPS-dependent degradation of IκBα is a key step in the activation of NFκB by TNFα where the ubiquitination of IκBα is driven by Skp1-Cul1-β-TrCP (SCF^β-TrCP^),^47^ a member of the CRL1 family E3 ligases whose assembly and disassembly are regulated by the CSN; hence, the survival signaling from NFκB is likely suppressed by impairment of IκBα ubiquitination due to defective Cullin deneddylation resulting from Cops8 deficiency. Our prior study detected decreases in myocardial F-box protein β-TrCP protein levels in Cops8^CKO^ mice,^7^ adding a reason to predict a reduction of SCF^β-TrCP^ ligase activities. Thus, Cops8 deficiency swings TNFR1 signaling towards the cell death direction.

Then, the next question is why necroptosis instead of apoptosis takes place. At least in the case of induction of necroptosis by death receptor activation, two prerequisites must be met in the cell. First, the formation of the so-called complex 2 containing RIPK1 and RIPK3 and second, the failure of caspase 8 to activate.^11^ Indeed, we observed that both prerequisites were met in the Cops8^CKO^ hearts. Not only were RIPK1, RIPK3, and MLKL protein levels markedly increased but also RIPK1-intereacted RIPK3 was significantly increased (**Figure 1**); and very importantly the cleaved form of caspase 8 as well as caspase 8 activity were substantially lower in the homozygous Cops8^CKO^ hearts compared with CTL hearts (**Figure 7**). It is very likely that this impairment of caspase 8 activation directly results from the loss of Cullin deneddylation because a prior study has established that Cul3-RBX1 mediated polyubiquitination of caspase 8 is required for further processing and activation of caspase 8 and the signaling of the extrinsic apoptotic pathway.^16^ Both neddylation and deneddylation of Cullins are required for proper functioning of CRLs; hence, the ubiquitination of caspase 8 by Cul3-RBX1 is very likely suppressed by Cops8 deficiency. Besides caspase 8 which is essential to the extrinsic pathway of apoptosis, as discussed below, the mitochondrial pathway is likely suppressed by increased BCL2 (**Figure 8**).^7^

We have previously observed a suppressed autophagic flux in Cops8^CKO^ mice. This could probably be due to impairment in autophagosome-lysosome fusion that occurs before impairment in the UPS degradation of a surrogate misfolded protein as well as CM necrosis become discernible.^8^ We propose dual impairment of both the UPS and the ALP plays an overall causative role in the CM necrosis that now proves to be necroptosis. This proposition now has support from two recent studies that collected evidence from cultured H9c2 cells suggesting a major contribution from impaired autophagy to the induction of necroptosis by TNFα.^48, 49^ According to these reports, RIPK1-RIPK3 interaction and necroptosis induced by the combined treatment with TNFα and z-VAD-fmk (a broad spectrum caspase inhibitor) were associated with suppression of autophagic flux,^48^ improving autophagic flux via mTORC1 inhibition suppressed the necroptosis in an autophagy- and transcription factor EB (TFEB; a master regulator of the ALP)-dependent manner,^48, 49^ and MPT does not to play a major role in the execution of necroptosis.^48^ This scenario starkly resembles what we have unveiled in the Cops8^CKO^ mouse myocardium. Hence, in the future it will be interesting and important to test whether the impaired autophagic flux has exacerbated activation of the RIPK1-RIPK3 necroptotic pathway in Cop8^CKO^ mice.

### Sustained Nrf2 activation and reductive stress contribute to the CM necroptosis

A surprising discovery of this study is that the sustained activation of Nrf2 in CMs promotes CM necroptosis and mouse premature death in the Cops8^CKO^ mice. Our prior transcriptome analysis has revealed a marked upregulation of Nrf2 target genes in Cops8^CKO^ hearts at both 2 and 3 weeks of age,^43^ indicative of Nrf2 activation by Cops8 deficiency. The sustained activation of Nrf2 is reflected further by increases in both pS40-Nrf2 (an active form of Nrf2) and total Nrf2 protein levels in Cops8^CKO^ mouse hearts at both 2 and 3 weeks of age (**Figure 8A ~ 8C**) and by increased proteins and mRNA expression of BCL2 (**Figure 7D, 7E**), a known Nrf2 target gene.^50^ Here the Nrf2 activation is probably triggered by increased oxidative stress resulting from impaired protein quality control (PQC) and is sustained by the defective inactivation of Nrf2. We have previously reported that Cops8 deficiency impairs the performance of both the UPS and the ALP, thereby impairing important cardiac PQC mechanisms.^4, 7, 8^ Impaired PQC is known to increase oxidative stress;^51^ indeed we detected increased myocardial levels of superoxide anions and protein carbonyls in mice with homozygous Cops8^CKO^ (**Figure 5**), compelling evidence of increased oxidative stress. As suggested by increased myocardial protein levels of both pS40-Nrf2 and total Nrf2 in Cops8^CKO^ mice at both 2 and 3 weeks of age (**Figure 8A~8C**), Cops8 deficiency likely impairs Nrf2 degradation. This is because Nrf2 degradation is mediated by the UPS and the responsible ubiquitin ligases are KEAP1-Cul3-Rbx1andβTrCP-Cul1-Rbx1, both belonging to the CRL family.^52, 53^ Cullin deneddylation by the CSN requires all 8 COPS subunits to form the holocomplex and is essential to the proper functioning of all CRLs;^54^ thus Cops8 deficiency impairs the catalytic dynamics of CRLs and thereby impairs Nrf2 degradation. Taken together, both reduced myocardial caspase 8 activity and upregulated BCL2 in Cops8^CKO^ mice can be explained by perturbation of cullin deneddylation by Cops8 deficiency and are likely responsible for suppression of the extrinsic and the intrinsic apoptosis pathways, respectively, allowing necroptosis to take place.

Previous reports have shown an important role of increased reactive oxygen species (ROS) in RIPK3-mediated necroptosis in cultured cells.^38, 55^ In TNFα induced necroptosis, the RIPK3-centered necrosome increases ROS production through stimulating aerobic metabolism and RIPK3 does so probably by activating key enzymes of metabolic pathways including glycogen phosphorylase (PYGL), glutamate-ammonia ligase (GLUL), glutamate dehydrogenase 1 (GLUD1),^38^ and more recently pyruvate dehydrogenase (PDH) which is a rate-limiting enzyme linking glycolysis to aerobic respiration.^56^ The increased ROS further promotes necrosome formation and yields cytotoxicity during necroptosis.^55^ As reflected by increased DHE staining of superoxide and the elevated levels of protein carbonyls in Cops8^CKO^ hearts (**Figure 5**), increases in ROS or oxidative stress are indeed associated with CM necroptosis in Cops8^CKO^ mice. Consistent with increased oxidative stress, Nrf2 and activated Nrf2, the master regulator of antioxidant and defensive responses, are markedly upregulated in Cops8^CKO^ hearts even before CM necrosis becomes discernible (**Figure 8A ~ 8C**). However, administration of a ROS scavenger NAC or CM-restricted overexpression of HMOX1 failed to reduce CM necrosis; on the contrary, these measures markedly increased CM necrosis or exacerbated mouse premature death in Cops8^CKO^ mice (**Figure 6**). Moreover, CM-restricted Nrf2 haploinsufficiency surprisingly delayed the premature death of Cops8^CKO^ mice (**Figure 8D**). These findings from the present study provide compelling evidence that sustained Nrf2 activation and resultant reductive stress, rather than ROS *per se*, contribute to the induction of CM necroptosis by Cops8^CKO^ in mice.

### Clinical implications

The discoveries of the present study have significant clinical implications. For example, first of all, inadequate cardiac PQC due to UPS malfunction and ALP impairment has been implicated in the progression from a large subset of heart disease to heart failure;^57, 58^ however, the mechanistic link between impaired PQC and heart failure has been obscure. The discoveries of the present study implicate that CM necroptosis could be one of the missing links, because cardiac PQC impairment is obviously the apical defect in Cops8^CKO^ mice. Accordingly, targeting the necroptotic pathway could potentially help alleviate the adverse outcome of cardiac PQC impairment. Second, a small molecule CSN inhibitor (CSN5i) that inhibits the cullin deneddylation activity of the CSN by specifically targeting Cops5 has shown great promise in anti-tumor effects in experimental studies.^18^ Hence, there is a good possibility for this compound to move into clinical trials for the treatment of cancer. CSN5i is expected to affect the degradation of a much smaller range of proteins than proteasome inhibitors would while being equally or even more effective in blocking cell cycle progression and causing cell death. The findings of the present study caution that cardiac function should be closely monitored should CSN5i or alike be moved into clinical trials. Lastly yet importantly, because of the wealth of accumulated evidence showing that Nrf2 is the major promotor of cellular defense against various pathological stresses in different organs, such as lungs, livers, kidneys, and the heart, Nrf2 has evolved to be an attractive drug target for the prevention or treatment of human diseases including heart failure.^59, 60^ However, a phase III clinical trial of bardoxolone methyl, an Nrf2 inducer, for the treatment of chronic renal disease associated with diabetes was terminated due to significantly increased incidence of heart failure.^61^ It is unclear whether the “dark” side of Nrf2 is linked to the magnitude of Nrf2 activation^62^ or simply due to off-target effects of the drug. Notably, a number of clinical trials at various phases on Nrf2 inducers for treating several other forms of disease (e.g., multiple sclerosis, cancers, pulmonary artery hypertension) are still ongoing; hence, elucidation of the mechanism governing the dark side of Nrf2 activation on the heart is absolutely warranted. To this end, the discovery of the present study that sustained Nrf2 activation and reductive stress promote CM necroptosis in a heart with impaired functioning of autophagy and the UPS may provide a previously unsuspected mechanism for the adverse cardiac effect of Nrf2 inducers.

## Conclusions

In conclusion, the present study has discovered that CM necrosis in Cops8^CKO^ mice belongs to necroptosis; the activation of the RIPK1-RIPK3 pathway, sustained Nrf2 activation, and reductive stress but not MPT mediate the CM necroptosis. Since the key processes mediating the CM necroptosis here can be traced back to impaired functioning of CRLs, we demonstrate here that Cops8/the CSN by virtue of cullin deneddylation suppresses necroptosis and plays a crucial role in shaping the mode of regulated cell death. The emerging model for Cops8 deficiency to cause CM necroptosis is illustrated in **Figure 8E**. In brief, loss of cullin deneddylation resulting from Cops8^CKO^ perturbs the catalytic dynamics of all CRLs, which in turn dysregulates the ubiquitination of a large subset of proteins and thereby impairs many cellular processes such as UPS-mediated protein degradation and autophagosome maturation, resulting in PQC impairment, increased proteotoxicity, and oxidative stress. As a result, CMs and possibly their non-CM neighbors increase the expression and secretion of TNFα. The autocrinal and paracrinal TNFα then bind TNFR1 on CMs and initiate TNFR1-mediated cell survival and/or death signaling. The survival signaling via NFκB activation is impaired because the ubiquitin-dependent degradation of IκBα is driven by a CRL type of E3 ligase (SCF^βTrCP^) but the latter does not function properly due to Cops8 deficiency; consequently, the cell death pathways via formation of the RIPK1- or RIPK1-RIPK3-centered complex 2 become inevitable. Since caspase 8 activation and processing also requires Cul3-mediated polyubiquitination,^16^ caspase 8 is disabled when cullin deneddylation is shut down; hence, the RIPK1-RIPK3 complex takes its course to necroptosis. Probably by upregulating anti-apoptotic factors such as BCL2 as well as causing reductive stress, sustained Nrf2 activation due to the impaired inactivation and degradation also helps steer the cell death mode to necroptosis, a more damaging form of cell death than apoptosis.

