## Supplemental Table and Figures for "The COP9 Signalosome Suppresses Cardiomyocyte Necroptosis"

**Supplementary Table S1**

**Supplementary Figures S1 and S2**

| **Supplementary Table S1. PCR primers used for genotyping** | | |
| --- | --- | --- |
| **PCR** | **Primer** | **Sequence (5'-3')** |
| *Cops8^flox^* | Forward | GACTCCACCACAAGAATGGTT |
|  | Reverse | GTAGGTGACCTTCAATGTGAC |
| *HMOX1* overexpression | Mutant Forward | TGCATCGCATTGTCTGAGTAGG |
|  | Wild-Type Forward | GGGGCGTGCTGAGCCAGACCTCCAT |
|  | Shared Reverse | TCCCGACAAAACCGAAAATCTGTGG |
| *Myh6-Cre^Tg^* | Transgene Forward | ATGACAGACAGATCCCTCCTATCTCC |
|  | Transgene Reverse | CTCATCACTCGTTGCATCGAC |
|  | Wild-Type Forward | CAGCTTCAGGAACAGCAGGTCC |
|  | Wild-Type Reverse | CATCAATCTCGCAGGTGTAGGACT |
| *Nrf2^flox^* | Forward | TCATGAGAGCTTCCCAGACTC |
|  | Reverse | CAGCCAGCTGCTTGTTTTC |
| *Ppif^-/-^* | Knockout Forward | GGCTGCTAAAGCGCATGCTCC |
|  | Wild-Type Forward | CTCTTCTGGGCAAGAATTGC |
|  | Shared Reverse | ATTGTGGTTGGTGAAGTCGCC |
| *RIPK3^-/-^* | Knockout Forward | CCAGAGGCCACTTGTGTAGCG |
|  | Wild-Type Forward | GCAGGCTCTGGTGACAAGATTCATGG |
|  | Shared Reverse | CGCTTTAGAAGCCTTCAGGTTGAC |

**A**

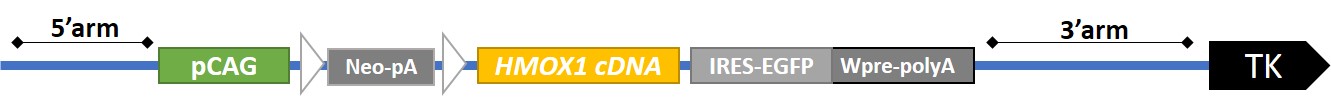

**B**

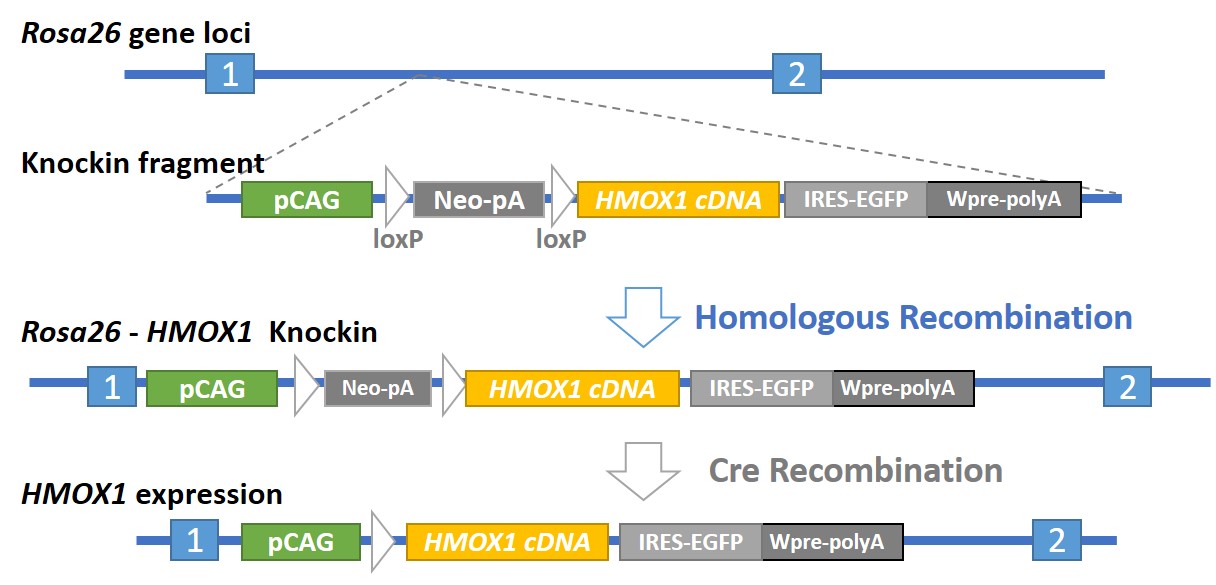

**Supplementary Figure S1**. An illustration of strategies for creation of R26-(CAG-LNL-HMOX1)1 mouse and the use of the mouse for conditional overexpression of HMOX1. **A**, A scheme of the vector used for targeting the conditional HMOX1 expression cassette into the Rosa 26 locus of mouse genome. **B**, Illustrations of strategies for the creation of R26-(CAG-LNL-HMOX1)1 mouse and the use of the mouse for conditional overexpression of HMOX1.

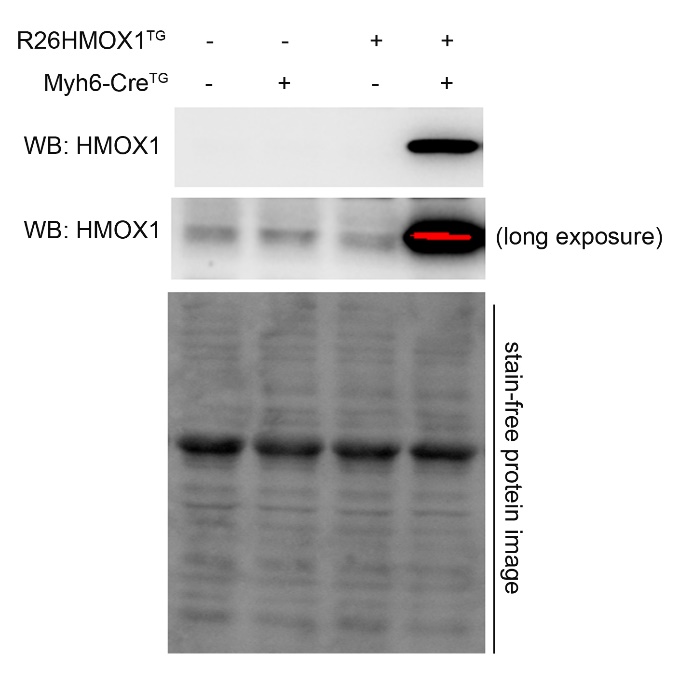

**Supplementary Figure S2**. Western blot analysis for myocardial expression of Heme Oxygenase 1 (HMOX1) in mice with the indicated genotypes. The stain-free protein image serve as loading control.
